# CTLA-4 checkpoint signalling modulates germinal centre dynamics and antibody avidity during *Borrelia burgdorferi* infection in mice

**DOI:** 10.64898/2026.10.30.763258

**Authors:** Varpu Mäkinen, Jukka Alinikula, Ruth Fair-Mäkelä, Pia Sundqvist, Marko Salmi, Pieta Mattila, Annukka Pietikäinen, Jukka Hytönen

## Abstract

*Borrelia burgdorferi*, the causative agent of Lyme borreliosis, persists in mammalian hosts despite robust activation of innate and adaptive immunity. A hallmark of murine infection is the production of low-avidity antibodies, yet the immunological mechanisms underlying this defective humoral response remain poorly understood.

Here, we used single-cell RNA sequencing of draining lymph node cells from Borrelia-infected mice at days 7 and 14 post-infection, comparing transcriptional profiles with those from mice immunised with heat-killed Borrelia or with a potent germinal centre-inducing protein immunisation. Borrelia infection induced a rapid germinal centre response, however, light zone B cells displayed diminished expression of co-stimulatory and T-B cell interaction molecules while maintaining core germinal centre identity markers, suggesting impaired positive selection. T follicular helper cells upregulated multiple inhibitory checkpoint receptor genes, most prominently Ctla4, alongside an active germinal centre engagement programme. Foxp3⁺ regulatory T cells upregulated genes associated with follicular-homing and suppressive function, including those implicated in CTLA-4-mediated trans-endocytosis.

To test whether inhibitory signalling contributes to the impaired humoral response, infected mice were treated with anti-CTLA-4 blocking antibody. CTLA-4 blockade significantly increased plasma cell numbers and enhanced both the magnitude and avidity of *Borrelia*- specific IgG responses, including the induction of complement-fixing IgG2a. These findings reveal that CTLA-4-dependent inhibitory signalling contributes to defective antibody quality during *Borrelia* infection, identifying a mechanism by which dysregulated T cell responses may facilitate impaired humoral control of a persistent spirochaetal pathogen.

**Authors summary:** *Borrelia burgdorferi*, the bacteria that cause Lyme borreliosis, can persist in the body for long periods despite triggering a strong immune response. One puzzling feature of this infection is that the antibodies produced are often of poor quality and do not bind the bacteria effectively.

To investigate why, we examined individual immune cells in lymph nodes of *Borrelia*-infected mice and compared them with cells from mice that received non-living *Borrelia* or a vaccine- like immunisation known to generate strong antibody responses. Although *Borrelia* infection rapidly triggered the formation of structures that normally help the immune system refine and improve antibodies, key communication signals between immune cells were reduced. At the same time, both helper and regulatory T cells showed increased activity of molecular pathways that act as brakes on immune responses. One such brake, controlled by a molecule called CTLA-4, appeared particularly important. When we blocked CTLA-4 during infection, mice produced more antibody-secreting cells and generated antibodies that had improved functional properties.

Our findings identify a mechanism that limits antibody quality during *Borrelia* infection and help explain how the bacteria can evade effective immune control and persist in the host.

## Introduction

*Borrelia burgdorferi* sensu lato spirochetes (hereafter *Borrelia*) cause Lyme borreliosis, the most common vector-borne disease in Northern Hemisphere (1). In natural reservoir host species, *Borrelia* infection is typically asymptomatic, demonstrating effective co-evolution of the bacterium and its hosts (2,3). Although *Borrelia* infection triggers both innate and adaptive immune responses, these responses fail to clear the infection, leading to bacterial persistence and consequently transmission to new hosts via ticks (2). In accidental hosts such as humans, untreated *Borrelia* infection can persist and disseminate to multiple tissues leading to more severe manifestations of the infection such as Lyme neuroborreliosis and Lyme arthritis (1).

Our recent study identified a potential mechanism suppressing T-B cell communication during *Borrelia* infection (4). We found that early-stage *Borrelia* infection induced upregulation of Suppressor of cytokine signalling 1 and 3 (*Socs1* and *Socs3*) in B cells, potentially impeding upregulation of genes related to T-B cell communication. *Borrelia* induces strong extrafollicular responses with sustained antibody production (3,5,6). However, the antibody production is IgM-biased and fails to generate long-lasting neutralizing immunity (3,7). In mice, B cells accumulate in lymph nodes, but germinal centre (GC) responses are impaired (8,9). Within the GC light zone (LZ), T follicular helper (Tfh) cells help select high-affinity B cell clones. Although the quality of T cell help has been widely studied, its contribution to poor antibody affinity maturation during *Borrelia* infection remains unclear (10,11). Available evidence indicates that although T cells, including Tfh cells, are effectively activated during *Borrelia* infection, their function might be modulated by inhibitory checkpoint pathways such as PD-1/PD-L1 and CTLA-4/B7.

In this study, we performed single-cell RNA sequencing (scRNA-seq) of lymph node lymphocytes at days 7 and 14 after *Borrelia* infection in mice, revealing upregulation of immune checkpoint receptors in Tfh cells. We reasoned that enhancing T cell help would normalize the immune response and improve antibody affinity maturation (12). To test this, we treated *Borrelia*-infected mice with anti-CTLA-4 and anti-LAG-3 antibodies. CTLA-4 blockade improved hallmarks of antibody-mediated immunity. Our findings suggest that expression of immune checkpoint receptors on Tfh cells, possibly together with Tfr cell activity, contributes to poor antibody affinity maturation during *Borrelia* infection.

## Results

### Borrelia infection induces excessive B cell cellularity in lymph nodes and GCs

To characterize the lymph node lymphocyte composition and investigate the molecular mechanisms underlying abnormal GC reactions, we collected draining popliteal lymph nodes from *Borrelia*-infected mice at days 7 and 14 post-infection (n=3 for both Bb7 and Bb14). Three control groups were included: mice immunised with heat-killed *Borrelia* (n=3 for both Bb_HK7 and Bb_HK14), mice immunised with ovalbumin in Complete Freund’s Adjuvant (n=3 for both OVA7 and OVA14), and PBS-injected mice as vehicle controls (n=3 for both PBS7 and PBS14) (Fig. 1A). Success of infection was confirmed by *Borrelia*-specific *flaB* PCR of urinary bladders. All mice receiving live *Borrelia* were positive at day 14, whereas those receiving heat-killed *Borrelia* remained negative (data not shown). All mice injected with either live or heat-killed *Borrelia* developed a *Borrelia*-specific antibody response, confirming seroconversion.

**Figure 1.**
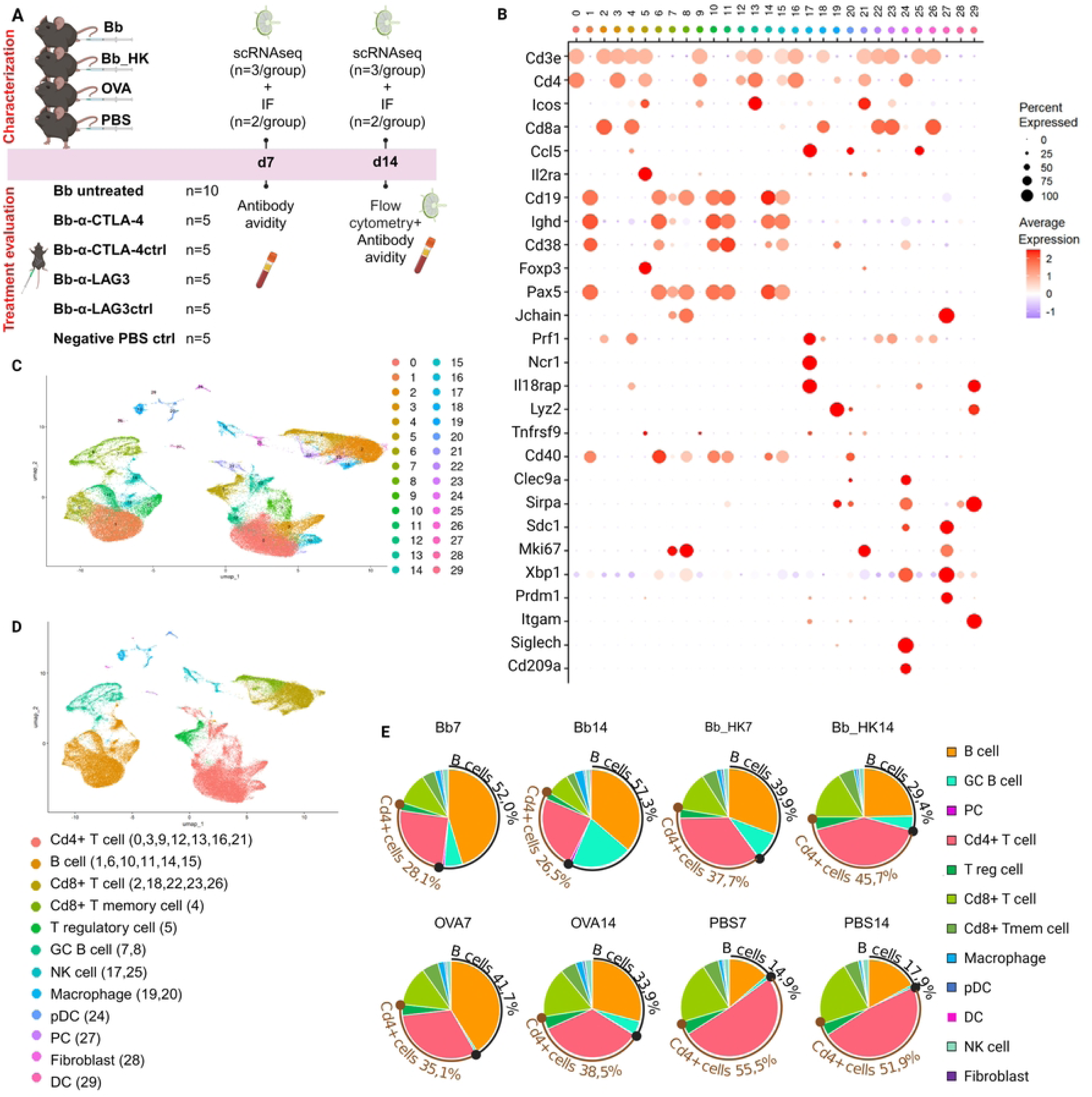
*Borrelia* infection induces excessive B cell cellularity in lymph nodes. (A) Experimental outline of single-cell RNA sequencing and immune checkpoint blockade treatments. (B) Expression of specific marker genes used to define cell clusters. (C) Clustering of all study groups together, based on the differentially expressed genes between clusters, resulted in 30 distinct clusters. (D) Identified major immune cell populations. Cluster numbers from (B-C) are indicated in brackets. Clusters were identified following data integration. (E) Relative proportions of cell types across study groups, indicating increased B cell abundance with infection. Abbreviations: GC: germinal center, NK: natural killer, pDC: plasmacytoid dendritic cell, PC: plasma cell, DC: dendritic cell, EFmem: extrafollicular memory, LZ: light zone, DZ: dark zone.

To explore the cellular and molecular basis of GC dysfunction during *Borrelia* infection, we performed scRNAseq on pooled lymph node samples (n=3 lymph nodes/pooled sample per study group). Clustering based on gene expression profiles identified 30 clusters, which were annotated into 12 major cell populations based on established marker genes: B cells, CD4+ T cells, CD8+ T cells, regulatory T cells (Treg), CD8+ memory T cells, GC B cells, plasma cells (PC), macrophages, plasmacytoid dendritic cells (pDC), dendritic cells (DC), natural killer (NK) cells, and fibroblasts (Fig. 1B-D, Supplementary 1).

A prominent B cell response was observed across study groups except the vehicle controls. At both days 7 and 14 after *Borrelia* infection, B lineage cells (including B cell, GC B cell and PC populations) constituted the largest lymphocyte fraction (Bb7: 52.0 % and Bb14: 57.3 %, Fig. 1E).

### B cell subclustering

To investigate the previously described abnormalities in T-B cell communication (4), we further characterized B cells and CD4+ T helper cells. Re-clustering of B lineage populations yielded nine subclusters, identified using established marker genes and transcription factor activity (Fig. 2A, Supplementary 2): resting, naive follicular, T cell-activated, interferon- activated extrafollicular, extrafollicular memory, light zone (LZ), dark zone 1 (DZ1), proliferating dark zone 2 (DZ2), post-GC B cells and PCs. One low-quality cluster and two T cell contaminated clusters were excluded from further analysis (Supplementary Fig. 1A-D).

**Figure 2.**
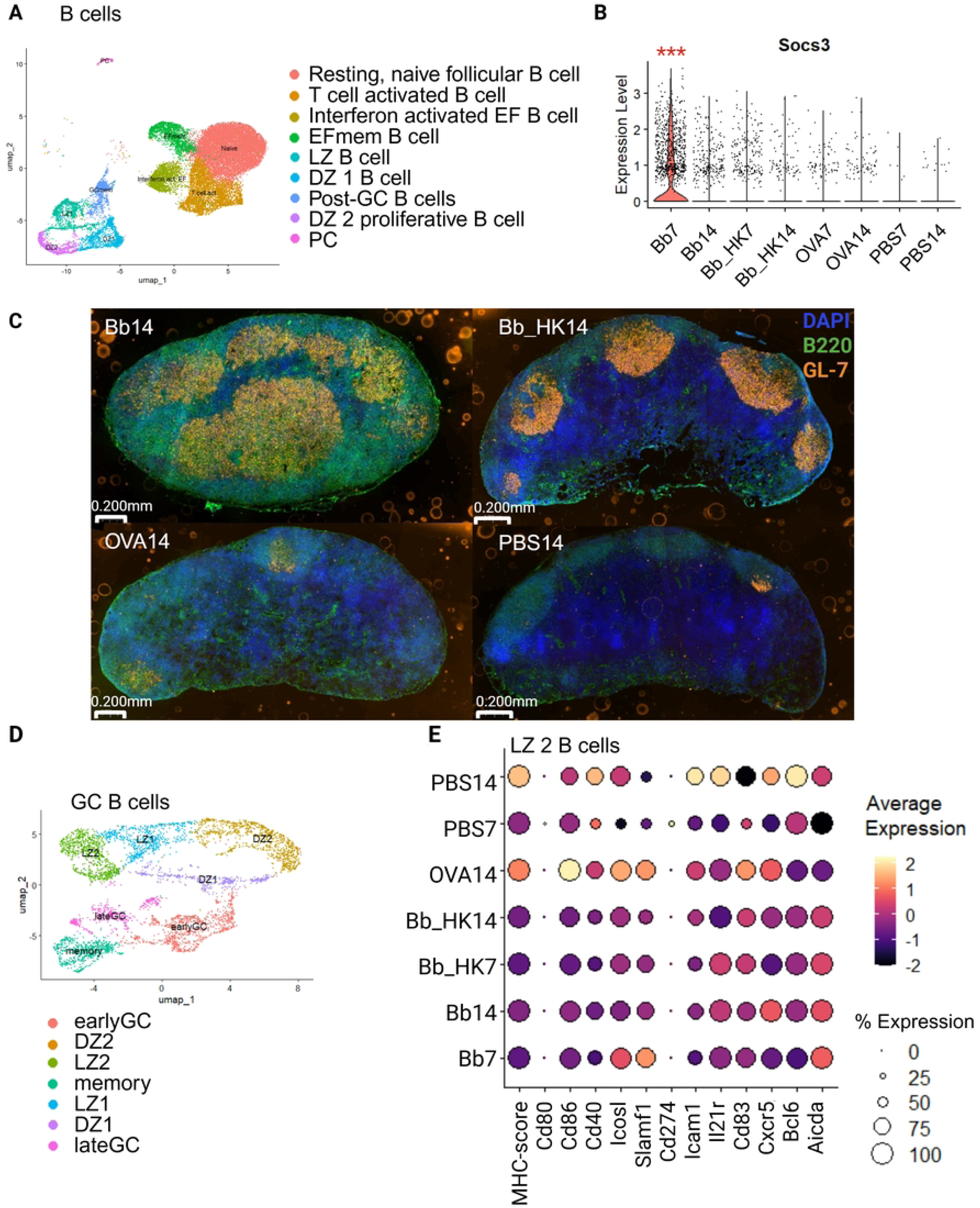
Altered B cell and GC B cell gene expression after *Borrelia* infection. (A) Re-clustering of B cells into subpopulations. (B) Expression levels of *Socs3* in activated B cell subsets (T cell activated B cell, Interferon activated EF B cell and EFmem B cell clusters analysed together) were significantly upregulated 7 days after *Borrelia* infection. (C) Immunofluorescence imaging of draining lymph nodes 14 days after *Borrelia* infection and control stimuli. DAPI (blue), B220 (B cells, green), and GL7 (germinal centers/activated B cells, orange) illustrating accumulated B cells after infection. (D) Re-clustered GC cells from the B cell subset. (E) Reduced expression of key Tfh-B cell interaction genes in LZ B cells during infection (MHC score: percentage of reads mapping to *H2* genes). Abbreviations: EFmem: extrafollicular memory, LZ: light zone, DZ: dark zone.

At day 14, GC B cells were enriched in *Borrelia*-infected mice compared with all other conditions (Bb7 11.1 %, Bb14 31.3 %, Bb_HK7 19.5 %, Bb_HK14 16.0 %, OVA7 0.7 %, OVA14 11.5 %, PBS7 2.5 %, PBS14 1.4 %, Fig. 1E). Consistent with previous findings (4), activated B cell subsets (T cell-activated, interferon-activated extrafollicular, and extrafollicular memory B cells, analysed collectively) showed significant upregulation of *Socs3* at day 7 after *Borrelia* infection, which may suggest reduced antigen presentation in early stages of the infection (Fig. 2B).

Immunofluorescence imaging of the draining lymph nodes at day 14 revealed striking differences in lymph node architecture across conditions (Fig. 2C). In *Borrelia*-infected mice, lymph nodes were enlarged with extensive B220+ B cell accumulation and widespread GL7+ staining beyond discrete follicular structures. In contrast, lymph nodes from mice immunised with heat-killed *Borrelia* maintained organised architecture, with defined B cell follicles and distinct compartmentalisation into cortex and paracortex despite the similar initial antigen load. OVA-immunised mice showed defined follicles and small GCs, while PBS controls exhibited minimal B cell accumulation and sparse GL-7 staining, consistent with the absence of antigenic stimulation (Fig. 2C). These findings are consistent with previously reported disruption of lymph node architecture during *Borrelia* infection (9,13).

### Impaired Tfh–B cell interaction in germinal centres

To assess T cell help during the GC reaction in *Borrelia* infection and to better define GC subpopulations, we re-clustered GC B cells identifying seven subclusters: early GC, DZ1, DZ2, LZ1, LZ2, late GC, and memory B cells (Fig. 2D, Supplementary 3). We then performed a comparative gene ontology (GO) analysis on the LZ1 and LZ2 B cell populations, the subsets most likely to interact with Tfh cells, using differentially expressed genes across experimental groups and time points (Supplementary Fig. 2). The analysis suggests that *Borrelia* infection is associated with reduced endocytosis, potentially leading to impaired antigen presentation (Bb14 vs. Bb_HK14; Supplementary Fig. 2A), as well as diminished T cell activation, which may reflect weakened T–B cell interaction (Bb14 vs. OVA14; Supplementary Fig. 2B). Within the *Borrelia* infection, LZ B cells at day 14 (Bb14 vs. Bb7) showed downregulation of genes involved in B cell activation (Supplementary Fig. 2C).

To further evaluate T cell help within the LZ, we examined expression of canonical Tfh-B cell communication genes in the LZ2 B cells, which showed the highest expression of LZ marker genes (Figure 2E). The OVA7 group was excluded from the analysis because an insufficient number of cells was recovered for reliable downstream analysis. Across all *Borrelia*-infected and heat-killed *Borrelia*-immunised groups, LZ2 B cells displayed lower expression of the co-stimulatory molecules *Cd80, Cd86*, and *Cd40*, together with a reduced MHC score (percentage of reads mapping to *H2* genes), relative to OVA-immunised mice at day 14. Importantly, we found no evidence that T-B cell interactions strengthened as the *Borrelia* infection progressed, contrary to expectations for an ongoing adaptive immune response. Within the *Borrelia*-infected group, LZ2 B cells showed a progressive decline in expression of *Icosl* and *Slamf1* from day 7 to day 14, although these trends did not reach statistical significance (Fig. 2E). ICOS–ICOSL interactions are critical for Tfh-mediated B cell selection (14) , while SLAMF1 mediates stable, long-lasting Tfh–B cell contacts required for sustained GC selection (15). In contrast, expression of *Cxcr5, Aicda*, and *Bcl6* increased or was maintained from day 7 to day 14, indicating that LZ B cells maintained an active GC programme during the later stages of infection. The dissociation of declining Tfh interaction molecules alongside sustained GC identity suggests that GC B cells in *Borrelia*-infected mice remain GC competent but progressively lose the capacity to engage Tfh cells for positive selection.

### Tfh cells display immune checkpoint receptor-high phenotype

To further characterize CD4+ T cell populations, we re-clustered these cells and identified eight subclusters based on their marker genes and transcription factor activity: naïve CD4+ T cells, three activated CD4+ T cell clusters (Act.1, Act.2, Act.3), InterferonI-stimulated T cells, Treg cells, IL21^high^ Tfh, and proliferating Tfh cells (Fig. 3A, Supplementary 4). One low-quality cluster was excluded (Supplementary Fig. 3A-B). At day 14, CD4+ T cells from *Borrelia*-infected mice showed significantly increased *Ctla4* expression relative to controls (Bb14 vs. Bb_HK14 p = 2.15e-51, Bb14 vs. OVA14 p = 9.45e-38, Fig. 3B).

**Figure 3.**
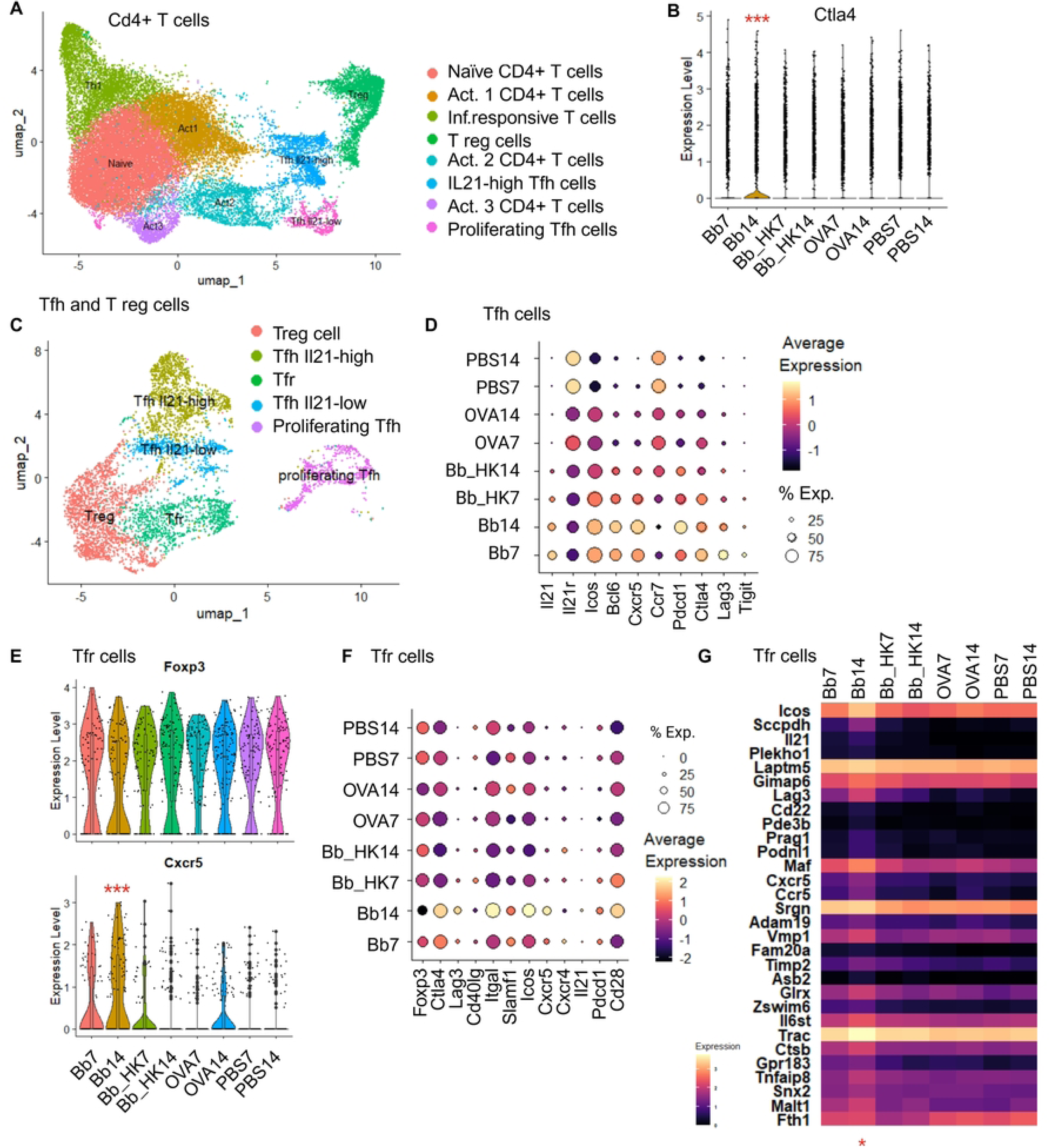
Gene expression profiles reveal expression of immune checkpoint receptors in T follicular helper cells and increased T follicular regulatory cell activity. (A) Re-clustering of CD4+ T cells into distinct subsets. (B) Expression of the gene encoding checkpoint molecule CTLA-4 was significantly upregulated 14 days after *Borrelia* infection. (C) Re-clustered Tfh and T reg cells from the Cd4+ T cell subset. (D) Tfh cells with active phenotype in GC response and with upregulation of immune checkpoint receptors during *Borrelia* infection. (E) *Cxcr5* expression in *Foxp3*+ Tfr population across study groups. (F-G) Relative expression of genes linked to migration into B cell follicles and intracellular trafficking/lysosomal activity in Tfr cells, indicating increased regulatory activity in GCs during *Borrelia* infection. * = adjusted P value < 0.01; ** = adjusted P value < 0.001; *** = adjusted P value < 0.0001. Abbreviations: Tfh: T follicular helper cell, Tfr: T follicular regulatory cells.

To examine its potential role in regulation of GC response, we re-clustered the combined populations of Tfh and Treg cells (Fig. 3C, Supplementary Fig. 4A-B, Supplementary 5), identifying five subclusters: Tfh IL21-high, Tfh IL21-low, proliferating Tfh, Treg, and Tfr. Tfh cells specifically from *Borrelia*-infected mice displayed a transcriptional program indicative of active GC engagement (Fig. 3D), with increased expression of *Cxcr5, Bcl6, Il21*, and *Icos* accompanied by reduced *Ccr7* expression from day 7 to day 14 consistent with migration from the T cell zone toward the GC. Despite the active GC programme, Tfh cells during *Borrelia* infection upregulated multiple inhibitory receptor genes, such as *Ctla4, Lag3*, *Tigit*, and *Pdcd1* (Fig. 3D). In contrast, Tfh cells of OVA-immunised mice showed minimal *Bcl6* and *Cxcr5* expression at these time points, likely reflecting the slower kinetics of the GC response following single protein immunisation.

Together, these data indicate that Tfh cells in *Borrelia*-infected mice acquire a GC-engaged transcriptional programme while simultaneously upregulating multiple immune checkpoint receptors.

### Borrelia infection promotes GC-homing suppression-competent phenotype in Tfr cells

Re-clustering of the combined Tfh and Treg populations identified a small population of Tfr cells (Fig. 3C). These cells expressed *Foxp3* across all study groups, confirming their regulatory identity (Fig. 3E). After *Borrelia* infection, Tfr cells distinctly upregulated *Cxcr5* compared with all other conditions (Fig. 3E-F), alongside increased expression of *Icos* (Fig. 3F), consistent with acquisition of a follicular-homing phenotype (Fig. 3F)(16).

Tfr cells from *Borrelia*-infected mice also acquired a cell-contact and suppressive programme. *Slamf1* was expressed at both day 7 and day 14, indicating early acquisition of T–B adhesion capacity (Fig. 3G). By day 14 post-infection, and in contrast to all other study groups, Bb14 Tfr expressed *Cd28* and *Itgal* (Fig. 3F). CD28 signalling is required for CTLA-4 expression and surface mobilisation in regulatory T cells (17) while LFA-1 (encoded by *Itgal*) stabilises the immunological synapse through ICAM-1 binding. These cells also showed increased expression of genes involved in endosomal trafficking and lysosomal activity (*Laptm5, Vmp1, Ctsb*) (Fig. 3G). By day 14 after *Borrelia* infection, the Tfr signature also included elevated expression of *Maf*, encoding the transcription factor c-Maf that promotes follicular T cell and regulator T cell function (18,19) and *Malt1*, required for Treg development (20), both highest in 14 days post-infection cells. Together, the co-expression of *Cd28*, *Itgal*, and *Ctla4* with endosomal and lysosomal genes indicates that 14 days post-infection, Tfr cells have assembled the molecular programme for CTLA-4-mediated trans-endocytosis (21) with synapse initiation, contact stabilisation, and co-stimulatory ligand capture and intracellular degradation.

Furthermore, a minor infection-specific increase in expression of *Gpr183*, encoding for EBI2 may suggest Tfr localization to B cell-T cell border. Although Tfr cells were not significantly enriched in number during *Borrelia* infection, their transcriptional profile at day 14 suggested a more mature, GC-homing phenotype, and suppression-competent phenotype distinct from all control conditions.

### CTLA-4 blockade improves antibody avidity against Borrelia

We first verified that CTLA-4 protein was localized within GCs during *Borrelia* infection (Fig. 4A). By comparison, CTLA-4 was not detected in GCs of OVA and heat-killed *Borrelia*-immunised mice (Supplementary Fig. 5A-B). We then tested whether blocking CTLA-4 and LAG-3 with antibodies could improve the antibody response against *Borrelia*. CTLA-4 blockade is known to induce Treg expansion via a feedback mechanism (22,23). Flow cytometric analysis of draining lymph nodes confirmed significant enrichment of Treg cells following anti-CTLA-4 treatment, indicating effective blockade (Supplementary Fig. 6). Consistent with elevated *Ctla4* transcript levels in scRNA-seq data, CD4+ T cells and Tfh cells expressed CTLA-4 protein as detected by flow cytometry (Fig. 4B). LAG-3 protein was low (Fig. 4B) as expected from low transcript level (Fig. 3E).

**Figure 4.**
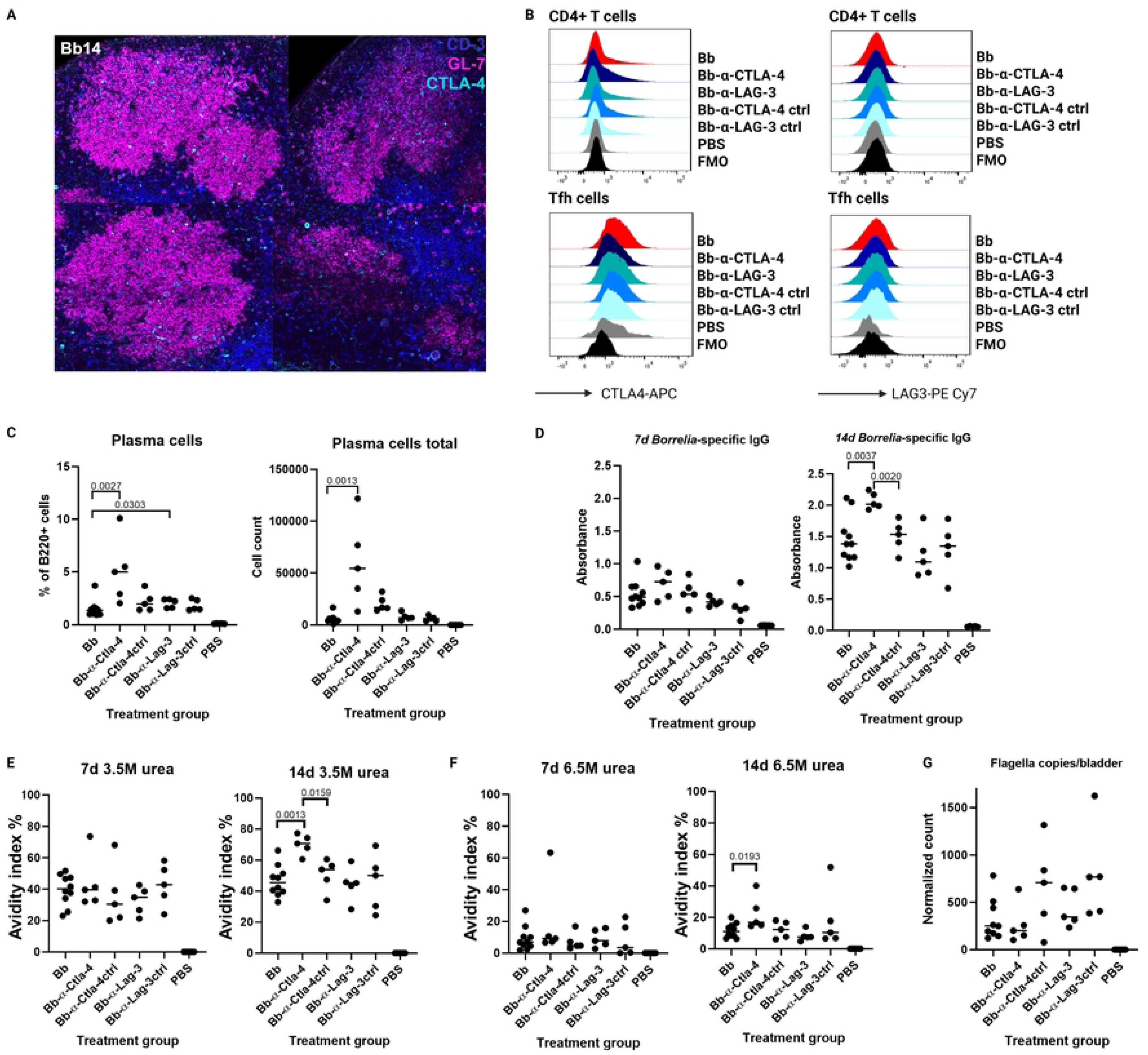
Blockade of the T cell checkpoint receptors CTLA-4 and LAG-3 in *Borrelia*-infected mice. (A) Immunofluorescence images showing CTLA-4 protein expression within GCs 14 days after *Borrelia* infection. CD3+ T cells (blue), CTLA-4 (cyan), and GL7 (germinal centers/activated B cells, magenta). (B) Mean fluorescence intensity (MFI) of CTLA-4 and LAG3 in CD4+ T cells and Tfh cells. (C) Frequency and number of CD138+B220low plasma cells after CTLA-4 or LAG-3 blockade. (D) *Borrelia*-specific IgG antibody concentrations at 7- and 14-days post-infection. Anti-CTLA-4-treated mice had significantly higher *Borrelia*-specific IgG concentrations than controls at day 14. Antibody avidity measurements demonstrating increased antibody strength after CTLA-4 blockade using 3.5M (E) and 6.5M (F) urea washes at 7- and 14-days post-infection. (G) Bacterial load in urinary bladders of from mice treated with CTLA-4 or LAG-3 blockade presented after Ct-based normalization to the mouse nidogen gene. CTLA-4 blockade showed decreasing trend that was not statistically significant. One datapoint in untreated group is out of range (not shown). P values < 0.05 were considered significant. Abbreviations: MFI: mean fluorescence intensity, FMO: fluorescence minus one.

CTLA-4 blockade increased CD138^+^B220^low^ plasma cell frequency compared with untreated mice (p = 0.0027) and isotype control (p = 0.0303) and this was reflected in total plasma cell numbers (p=0.00130 vs untreated; Fig. 4C). Importantly, by day 14 post-infection, anti-CTLA-4-treated mice had significantly higher *Borrelia*-specific IgG concentrations than untreated (p=0.0037) and isotype-control mice (p=0.0020; Fig. 4D).

To assess antibody quality, we calculated avidity indices from *Borrelia*-specific ELISA by comparing absorbances after urea vs. PBS washes. CTLA-4 blockade led to significantly increased avidity by day 14 under 3.5M urea compared with both untreated (p=0.0013) and isotype control mice (p=0.0159; Fig. 4E). This increase remained significant after 6.5M urea wash compared with untreated mice (p=0.0193; Fig. 4F). As expected, neither IgG concentrations nor antibody avidity increased significantly by day 7 (Fig. 4D-F). LAG-3 blockade had no effect on IgG concentrations or avidity, (Fig 4D–F).

To assess whether the increased avidity translated to better control of bacterial dissemination, we quantified bacterial load by *Borrelia*-specific *flaB* PCR from bladder tissue. No significant difference was observed between treatment groups, though anti-CTLA-4-treated mice showed a trend toward reduced bacterial burden compared with untreated and isotype-control groups (Fig. 4G). One outlier in the untreated group was excluded from analysis.

### Borrelia infection induces IgG subclass skewing that is partially rescued by CTLA-4 blockade

We analysed immunoglobulin heavy chain expression in combined GC B cells and plasma cells from the scRNA-seq data to characterise class-switching patterns during *Borrelia* infection. Live *Borrelia* infection induced a pronounced shift in isotype usage by day 14 where *Ighg2b* was the dominant class-switched transcript (Bb14 vs. Bb_HK14 p=7.38e-20; Bb14 vs. OVA14 p=1.08e-25; Fig. 5A). *Ighg3* was also upregulated compared with OVA14 (p = 4.28e-3), while I *Ighg1* was markedly downregulated in *Borrelia*-infected mice compared with OVA-immunised mice (p=3.75e-15). *Ighg2a* probe was absent from the scRNA-seq probe panel.

**Figure 5.**
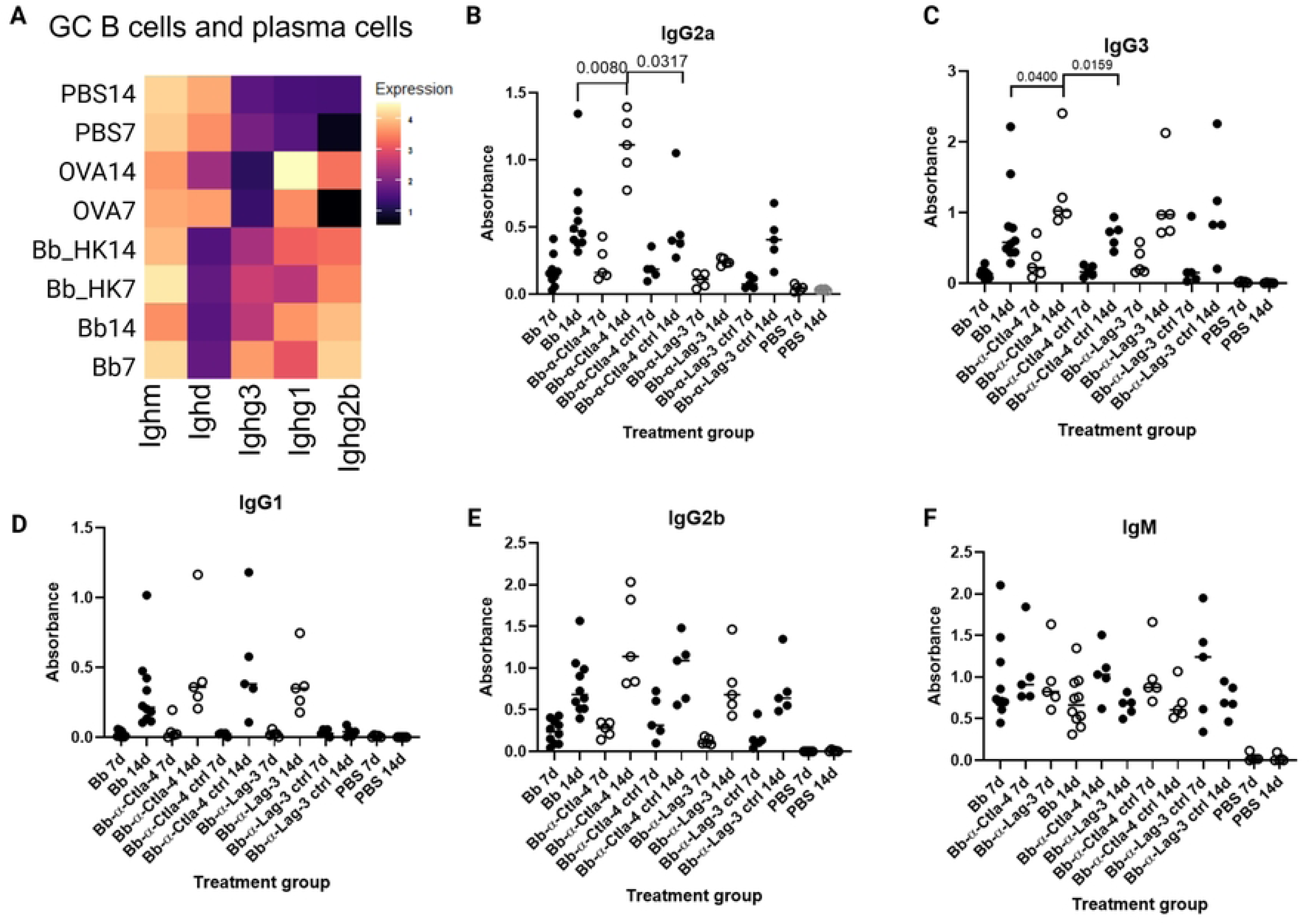
*Borrelia*-induced antibody isotype switching is affected by the CTLA-4 and LAG-3 checkpoint blockade. (A) Isotype-specific immunoglobulin gene expression across the study groups. (B-F) Serum antibody concentrations for different isotypes over time. CTLA-4 blockade increases IgG2a and IgG3 responses, while IgG1, IgG2b, and IgM remain unchanged, demonstrating selective effects on antibody class switching during *Borrelia* infection.

To assess whether immune checkpoint signalling influences subclass distribution at the protein level, we measured *Borrelia*-specific IgG subclass concentrations by ELISA. CTLA-4 blockade significantly increased IgG2a concentrations at day 14 compared with untreated (p = 0.0080) and isotype-control treated mice (p = 0.0317; Fig. 5B), and similarly increased IgG3 concentrations (p = 0.0400 and p = 0.0159, respectively; Fig. 5C). No significant changes in were observed IgG1, IgG2b, or IgM (Fig. 5D-F) indicating that CTLA-4 blockade selectively enhanced specific complement-fixing subclasses without altering the dominant IgG2b response. No significant differences in any subclass were observed at day 7. LAG-3 blockade had no effect on IgG subclass distribution at either time point (Fig 5B-F).

Together, these data indicate that *Borrelia* infection drives an IgG2b-dominant class-switched response in mice, and that CTLA-4 blockade broadens this response to include robust IgG2a production, the IgG subclass with the strongest complement activation capacity and thus the greatest potential to contribute to complement-dependent *Borrelia* clearance (24).

## Discussion

Despite extensive research on immunology of *Borrelia* infection, the mechanisms regulating GC responses remain incompletely understood. Here, we investigated the role of immune checkpoint signalling, particularly CTLA-4, in modulating the GC dynamics during experimental *Borrelia* infection. Although *Borrelia* infection induces Tfh cell differentiation and initiates GC reactions, GC organization and function are not sustained, limiting the production of long-lived, high avidity antibody-producing plasma cells (10). Our findings extend previous work on T-dependent immunity in *Borrelia* infection, which has largely focused on the loss of T cell help or co-stimulatory pathways (10,25), by demonstrating that CTLA-4-mediated signalling modulates GC dynamics and antibody quality. A previous study examining *Borrelia* infection in PD-1 knockout mice reported no effect on bacterial burden or serum *Borrelia* specific IgG antibody concentrations (11), suggesting pathway-specific roles for individual checkpoint receptors. We speculate that persistent antigen exposure from viable spirochetes within draining lymph nodes (9,26) may alter the local immune microenvironment and promote checkpoint receptor expression, a mechanism absent during exposure to heat-killed *Borrelia*.

Gene expression analysis of GC LZ B cells showed diminished expression of molecules involved in T-cell help while maintaining core germinal centre programs. Although not statistically significant, *Icosl* and *Slamf1* showed a trend towards downregulation from day 7 to day 14 in *Borrelia*-infected mice, while genes associated with ongoing GC activity (*Cxcr5*, *Bcl6*, *Aicda*) remained expressed. Reduced ICOS-ICOSL interaction could impair survival and selection of high-affinity B cell clones (14), while diminished SLAMF1 expression may compromise stable Tfh-B cell contacts essential for sustained GC selection (15). This pattern suggests that B cell remain within the GC programme but receive progressively less Tfh-derived signals, potentially due to the inhibitory follicular T cell phenotype described below.

Consistent with previous reports (10,11,25), our scRNA-seq data indicate that Tfh cells acquire an inhibitory receptor-high phenotype during *Borrelia* infection, with upregulation of *Ctla4, Lag3, Tigit and Pdcd1*. On Tfh cells CTLA-4 can limit co-stimulation through cell-intrinsic competition with CD28 for CD80/CD86 binding (27,28). Despite transcriptional upregulation of *Lag3*, LAG-3 protein expression on Tfh cells was minimal and LAG-3 blockade had no detectable effect on antibody responses, suggesting that CTLA-4, rather than LAG-3 or PD-1, is the functionally dominant checkpoint receptor in this context.

Our data further implicate Tfr cells in GC dysfunction during *Borrelia* infection (29,30). Although Tfr cell numbers were not increased during infection, their transcriptional profile at day 14 was markedly distinct from all other conditions. Tfr cells from *Borrelia*-infected mice acquired transcriptional programme consistent with suppressive competence from adoption of follicular phenotype, suppressive phenotype and adhesion molecules for sequential immune synapse initiation, stabilization, costimulatory ligand capture and endocytosis. While these transcriptomic findings cannot confirm functional trans-endocytosis, the coordinated expression of the complete molecular programme and its specificity to *Borrelia*-infected mice at day 14 represents a mechanism by which even small numbers of Tfr cells can exert potent suppressive effects within the GC (31).

The role of Tfr cells remains nuanced. Tfr cells did not express *Bcl6*, suggesting their maturation is in early stages at day 14. Tfr-mediated suppression can occur at the T-B border without full GC entry (28,32), where Tfr cells may primarily inhibit cytokine signalling rather than directly altering *Cxcr5* or *Bcl6* expression in Tfh cells (29,33). Consistent with this, *Il21* expression decreased from day 7 to day 14, indicating reduced helper capacity and *Gpr183* expression was slightly elevated in Tfr cells specifically during live *Borrelia* infection. The enrichment of genes associated with endosomal trafficking in Tfr cells is consistent with active CTLA-4-mediated trans-endocytosis. Thus, the combined effect of immune checkpoint receptor upregulation on Tfh cells and Tfr-mediated inhibition may jointly contribute to impaired affinity maturation.

CTLA-4 blockade significantly increased plasma cell numbers, *Borrelia*-specific IgG concentrations, and antibody avidity, supporting the conclusion that checkpoint signalling limits GC output and antibody quality during infection. CTLA blockade also broadened the class-switched response to include robust IgG2a production, the IgG subclass with the strongest complement-activating capacity (24), which is particularly relevant as complement-mediated clearance of *Borrelia* is heavily reliant on antibody-induced activation of the classical pathway (34–36). Despite these qualitative improvements, bacterial burden was not significantly reduced, suggesting that enhanced antibody maturation alone may be insufficient for bacterial clearance at this time point. Notably, the isotype control antibody influenced Treg and plasma cell populations, possibly reflecting a non-specific immune activation to a foreign antibody. While the anti-CTLA-4 effects remained significant compared with both untreated and isotype controls for IgG concentrations and avidity, this observation warrants consideration when interpreting plasma cell data.

Our results are consistent with Hastey et al. who showed that persistent IgM contributes to controlling bacterial burden but has limited impact on dissemination, highlighting the insufficient protective capacity of non-class-switched responses (37). CTLA-4 blockade enhanced class-switched IgG responses and antibody avidity, demonstrating that improving T cell help can partially overcome the impaired humoral response during *Borrelia* infection. However, bacterial burdens remained unchanged which raises important questions. Either the improved antibodies are still insufficient, the time frame is too short, or antibody-mediated clearance is not the primary mechanism of bacterial control. These findings align with recent evidence showing that Tfh responses are effectively induced but functionally constrained during *Borrelia* infection (38), and may have relevance for human Lyme borreliosis, where prolonged IgM responses and delayed antibody maturation have been reported (39,40).

A few limitations of this study should be considered. Samples were pooled for scRNA-seq (n = 1 per study group), precluding assessment of inter-individual variability. Also, some transcriptional differences did not reach statistical significance, likely reflecting limited sample sizes and cell numbers. The use of laboratory *Borrelia* strain N40 and needle inoculation does not fully recapitulate the immune modulation associated with natural tick-borne transmission (41,42), and the high inoculation dose may independently affect GC responses through excessive TLR2-mediated innate activation by *Borrelia* lipoproteins. However, similar high-dose inoculation models are well established (43–45), and our results align with previously described immune response patterns (9). Systemic anti-CTLA-4 treatment blocks CTLA-4 on all expressing cell types, including Tfh cells, Tfr cells, and conventional Tregs, and the relative contribution of each population to the observed effects cannot be determined from these experiments.

Collectively, our data support a model in which persistent antigen exposure during *Borrelia* infection induces a checkpoint receptor-high state in Tfh cells, coupled with acquisition of suppression competent Tfr cells. These mechanisms may act as brakes on GC responses, leading to impaired T cell help, reduced B cell selection, low-avidity antibody production, and skewed isotype production. CTLA-4 blockade partially reverses these defects, enhancing antibody quantity, avidity, and subclass breadth. Future studies combining cell type-specific checkpoint modulation with longer-term infection models will be instrumental to determine whether enhancing antibody quality translates into improved bacterial clearance.

## Materials and methods

### Borrelia preparation

*Borrelia burgdorferi* sensu stricto strain N40 was cultured in Barbour-Stoenner-Kelly II (BSK II) medium at +33 °C until mid-logarithmic phase. After washing bacteria twice with phosphate-buffered saline (PBS), cells were counted and concentrated to 50 × 10^6^ bacteria/ml in PBS. Heat-killed bacteria were inactivated in +56 °C for two hours. Successful inactivation was confirmed by the loss of motility under microscopic evaluation and six-week cultivation in BSK II medium, with no growth observed.

### Mouse experiments/study groups

Ten female mice (C3H/HeN, 4-5 weeks old, Inotiv, the Netherlands) were infected by administering 20 µl of the *Borrelia* suspension (10^6^ bacteria/injection; Bb7/Bb14) subcutaneously into the metatarsus of left hind leg (Microfine Demi 0.3 ml, BD #324826) and ten mice were injected with heat-killed *Borrelia* (Bb_HK7/Bb_HK14) in a same concentration (Fig. 1A). Ten mice were injected with Ovalbumin (Invivogen, vac-pova, 0.1 µg/µl) supplemented with Complete Freund’s Adjuvant (OVA7/OVA14, Sigma, 20% of the volume fraction). As a vehicle control, PBS (20 µl) was injected into nine mice (PBS7/PBS14). The mice were held under specific pathogen–free conditions at the Central Animal Laboratory of the University of Turku. All animal experiments were approved by the Ethical Committee for Animal Experimentation in Finland. Experiments were performed according to the Finnish Act on Animal Experimentation (497/2013) in compliance with the 3R-principle with the appropriate animal license (22423/2023 and 31189/2025).

After seven days, five mice from each group (four from PBS group) were euthanised, and the remaining mice were killed 14 days post-infection. Popliteal lymph nodes, draining the infection site, were harvested from three mice and combined per group for scRNAseq. Additionally, popliteal lymph nodes from two other mice per group and one from the PBS group were collected for histological analysis and embedded in Tissue-Tek optimal cutting temperature compound (O.C.T., Sakura #4583) in a pre-determined orientation to produce vertical sections of the tissue in relation to the hilar blood vessels. Samples were snap-frozen on dry ice, stored at -80 °C and used for cutting cryosections. Urinary bladders were collected for *Borrelia* PCR. Blood samples were collected with cardiac puncture, left to coagulate 30 minutes at room temperature (RT) and centrifuged 1500g for ten minutes after which sera were collected and stored at -80°C.

### Single-cell RNA-sequencing

Lymph nodes (n = 3/group) from each group were pooled into one sample. Single-cell suspension was produced by mechanically homogenising the pooled popliteal lymph nodes with a custom-made metal cell strainer to detach lymphocytes from the stroma. Cells were washed twice with PBS with 2 % fetal calf serum (FCS) buffer and filtered through 77 µm silk. Cells were counted with Cellometer Auto2000 (Revvity) and subsequently fixed 24 hours with Chromium Next GEM Single Cell Fixed RNA Sample Preparation Kit (10X Genomics) according to the protocol (10 X Fixation of Cells & Nuclei for Chromium Fixed RNA Profiling) and stored at -80 °C. Samples were processed with the Single Cell Gene Expression Flex (10X Genomics) assay in Single Cell Omics Facility of Turku Bioscience (Turku, Finland). Chromium mouse transcriptome probe set version 1.0.1 mm10-2020 was used. Single-cell libraries were sequenced using Illumina Novaseq6000 Sequencing System in the Finnish Functional Genomics Center of Turku Bioscience (Turku, Finland).

### Bioinformatic analysis

The raw data were pre-processed with the 10X Genomics Cell Ranger package version 8.0.1. Bioinformatic analysis was performed with R (version 4.4.1 2024-06-14) and RStudio (version 2024.9.1.394). For data analysis, Seurat – R tool kit (5.3.0) was used (46).

Each pooled sample group was subjected to quality control analysis separately. Cells with mitochondrial RNA content less than 10 % and between 400 and 7000 detected genes were retained for downstream analysis from Bb7, Bb14, Bb_HK7, Bb_HK14, OVA7 and OVA14 groups; for the PBS group, cells between 400 and 6000 detected genes were retained. Datasets from all experimental groups and time points were integrated into a single dataset prior to dimensionality reduction and clustering.

Principal component analysis was performed on normalized highly variable genes, and the first 40 principal components were used for UMAP visualization and nearest-neighbour graph construction. Clustering was performed with resolution 0.5. Cell clusters were identified with the minimum percentage argument set to 0.4, and the threshold for log2 fold change set to 0.5. For subcluster analyses, dimensionality reduction and nearest-neighbour graph were computed using 30 principal components. Clustering was performed using resolution 0.4 (*Cd4*+ T cells) or 0.3 (B cells, GC B cells, Tfh+Treg cells). Differentially expressed genes between the cell clusters and study groups were calculated using a Wilcoxon rank sum test. To support cell annotation, transcription factor activity was inferred using the DoRothEA (version 1.18.0) regulatory network resource and the univariate linear model (ULM) method implemented in decoupleR (version 2.12.0). Predicted transcription factor activities were used as complementary evidence with marker gene expression to improve cluster annotation.

Gene Ontology (GO) enrichment analysis was performed using clusterProfiler (version 4.14.6). Gene symbols were converted to Entrez Gene IDs using org.Mm.eg.db (version 3.20.0). Biological Process (BP) enrichment was made with Benjamini–Hochberg correction for multiple testing and an adjusted p-value cutoff of 0.05. Clustering strategy can be found in Supplementary Fig. 7. Cluster and subcluster analyses for each study group are presented with original cell counts in Supplementary Fig. 8.

### Immunofluorescence analysis

Frozen lymph node sections (8 µm; n=2 per group, PBS14 n=1) were fixed in ice-cold acetone 3 min and stained for immunofluorescence as described previously (4). DAPI (Invitrogen, conc. = 1 µg/ml) was used to visualize nuclei, B cells were identified with using Alexa Fluor® 488-conjugated monoclonal antibody against B220 (BioLegend, cat. #103225, clone = RA3-6B2, conc. = 10 µg/ml), and GC/activated B cells using Alexa Fluor® 647-conjugated monoclonal antibody against GL7 (Biolegend, cat. #144606, clone = GL7, conc. = 10 µg/ml). To detect CTLA-4, samples were stained with Rabbit monoclonal CTLA-4 antibody (E5S7E, Cell signaling technology, cat, #50490S, conc. = 10 µg/ml), and Goat anti-Rabbit IgG (H+L), Alexa Fluor™ 546 (Invitrogen, cat. #A11035, conc. = 5 µg/ml) was used as the secondary antibody. T cells were stained with Alexa Fluor® 488 anti-mouse CD3 Antibody (Biolegend, cat. #100210, clone = 17A2, conc. = 5 µg/ml).

### Blockade of immune checkpoint receptors CTLA-4 and LAG-3

Thirty mice were infected with *Borrelia* as described above. PBS, as a negative control, was injected similarly into five mice (Fig. 1A). Five *Borrelia* infected mice were treated with InVivoMAb anti-mouse CTLA-4 (CD152) (Clone:9H10, cat. BE0131, BioXcell) and five mice with InVivoMAb anti-mouse LAG-3 (Clone:C9B7W, cat. BE0174, BioXcell). Corresponding isotype control antibodies, InVivoMAb polyclonal Syrian hamster IgG (cat. BE0087, BioXcell) for the CTLA-4 and Purified Rat IgG1, κ Isotype Ctrl Antibody (cat. 400452, GoInVivo™) for the LAG-3, were administered as controls to five mice each. Antibody treatments were administered intraperitoneally at a dose of 200 µg per injection on infection days 0, 2, 4, 6, 8, 10, and 12. Untreated infected (n=10) and untreated PBS injected (n=5) mice served as controls. Blood samples were collected from the tail vein at 7 days post-infection and by cardiac puncture at 14 days post-infection. Samples were processed as described above and the avidity of *Borrelia*-specific antibodies was assessed. Popliteal lymph nodes were harvested for the flow cytometry analysis, and urinary bladders were collected to investigate the *Borrelia* load.

### Determination of Borrelia load in mouse tissues using semi-quantitative PCR

Urinary bladders were digested with ProteinaseK (4 mg/ml, 1 h at 56°C), and DNA was extracted using the NUCLISENS® easyMAG® system (elution 55 µl). *Borrelia* DNA was detected by PowerUp™ SYBR™ Green-based (Thermo Fisher #A25742) real-time PCR targeting the flagellin B gene (0.5 µM Fl7 5’-GCATTTTCAATTTTAGCAAGTGATG-3’ and 0.5 µM Fl59 5’-TTTCAGGGTCTCAGGCGTCTT-3’). Reactions (20 µl) contained 2 µl DNA.

To consider the differences in host tissue size and the yield of DNA extraction, bacterial copy numbers were normalized to the mouse nidogen gene (47) (0.5 µM MnidoF 5’-CCAGCCACAGAATACCATCC-3’ and 0.5 µM MnidoR 5’-GGACATACTCTGCTGCCATC-3’) using Ct-based normalization. Nidogen qPCR used 1 µl DNA in 25 µl reactions. Cycling conditions on the LightCycler®480 were: 50°C for 2 min, 95°C for 2 min, then 50 (*Borrelia*) or 40 cycles (nidogen) of 95°C for 15 s, 58°C (*Borrelia*) 57°C (nidogen) for 15 s, and 72°C for 1 min, with a melt curve analysis.

### Flow cytometry

Single-cell suspension was produced by mechanically homogenising each lymph node with 70 μm Falcon™ Cell Strainers (Fisher Scientific) to detach lymphocytes from the stroma. Cells were washed twice with PBS with 2 % fetal calf serum (FCS) and 0.01% sodium azide and filtered through 77 µm silk to remove cell aggregates.

Cells were stained with Fixable Viability Dye eFluor 780 (eBioscience, 65-0865-14, 1:1000) to identify live cells. To block the unspecific binding to Fc receptors, cells were incubated with an anti-CD16/32 antibody (BioXCell, BE0307, 1:100). The cells were stained with fluorescently labelled antibodies against T and B cell surface markers (Supplementary 6A). In fluorescence minus one (FMO) control the staining antibody was omitted. Cells were fixed overnight at +4° C and permeabilized with eBioscience Foxp3/Transcription factor staining buffer set (Thermo Fisher eBioscience #00-5523-00), following incubation with directly conjugated antibodies to detect the intracellular antigens (Supplementary 6A). Cells were analysed with a FACSymphony A5 SE flow cytometer with BD FACSDiva software (v9.6, BD Biosciences) and data were analysed with the FlowJo v10.10.0 Software (BD Life Sciences, gating strategy in Supplementary Fig. 9).

### Borrelia-specific antibody detection and antibody avidity index

*Borrelia*-specific IgM, IgG, and IgG subclass antibodies were measured using the Anti-*Borrelia* plus VlsE ELISA (IgG) kit (EUROIMMUN # EI 2132-9601-2 G) with mouse-specific conjugates (Supplementary 6B), following the manufacturer’s instructions with modifications. Samples were diluted (Supplementary 6B), added to microplate wells, incubated for 30 min at RT, and washed three times. For total *Borrelia*-specific IgG, enzyme-labelled secondary antibody was added and incubated for 30 min. For IgM and IgG subclasses, biotin-conjugated antibodies (50 µl) were added and incubated for 60 min, followed by washing and incubation with streptavidin-HRP (50 µl) for 60 min. TMB substrate was added, reactions were stopped with sulfuric acid, and absorbance was measured at 450 nm using a Victor Nivo Multimode Plate Reader (PerkinElmer).

To measure *Borrelia*-specific IgG antibody avidity, serum samples were diluted 1:200 and processed according to the respective protocols described above, with the following modifications. Each diluted sample was added to wells in three identical microplates. After the sample incubation and washing steps, one plate was treated with 100 µl of 3.5 M urea, a second with 100 µl of 6.5 M urea, and a third with 100 µl of PBS as a control. Plates were incubated for 15 min at RT, followed by washing as described in the standard protocol. The assay was then continued with conjugate incubation and subsequent steps as described above. The avidity index was calculated as the ratio of the absorbance value obtained after urea treatment to the corresponding absorbance value of the PBS-treated control and expressed as a percentage.

### Statistics

Normality of datasets was evaluated using the Shapiro–Wilk test. Significance between the cell numbers or antibody levels of treatment groups was evaluated either with an unpaired T-test or Mann-Whitney test depending on the normality of the data. The Wilcoxon rank-sum test was used to identify the differentially expressed genes and to compare the differences in the expression of genes of interest between the study groups with R software. The significance of differentially expressed genes was evaluated by adjusted p-value, based on Bonferroni correction. P value <0.05 (including adjusted values) was considered as statistically significant.

## Data availability

All scRNAseq data can be found from the GEO database with accession number GSE328572(48).

## Acknowledgements

The authors wish to acknowledge CSC – IT Center for Science, Finland, for computational resources. We thank Finnish Functional Genomics Centre supported by University of Turku, Åbo Akademi University, and Biocenter Finland, and Single Cell Omics Facility, Advanced Imaging Core, and Cytometry Core of Turku Bioscience. The authors would like to thank Etta-Liisa Virtomaa, Ann Sofie Wierda, and Diogo Cunha for the assistance. Figures of this article were created with Biorender.com.

## Author Contributions

Conceptualization: V.M., J.H, J.A., A.P., M.S., R.F.-M. Methodology: V.M., R.F.-M., P.S., A.P., J.A. Investigation: V.M., J.A., R.F.-M., J.H., A.P. Formal analysis: V.M., J.A., R.F.-M. Validation: V.M., J.A. Visualization: V.M., R.F.-M. Funding acquisition: J.H., A.P., J.A., Project administration: J.H. Supervision: J.H., J.A., A.P. Resources: J.H., J.A., P.M., M.S. Writing-original draft: V.M. Writing-review and editing: J.A., J.H., A.P., R.F.-M., P.S., P.M., M.S.

